# A Wnt-BMP4 signalling axis induces MSX and NOTCH proteins and promotes growth suppression and differentiation in neuroblastoma

**DOI:** 10.1101/2020.02.07.938654

**Authors:** Marianna Szemes, Zsombor Melegh, Jacob Bellamy, Madhu Kollareddy, Daniel Catchpoole, Karim Malik

## Abstract

The Wnt and bone morphogenetic protein (BMP) signalling pathways are known to be crucial in the development of neural crest lineages, including the sympathetic nervous system. Surprisingly, their role in paediatric neuroblastoma, the prototypic tumour arising from this lineage, remains relatively uncharacterised. We previously demonstrated that Wnt/β-catenin signalling can have cell-type specific effects on neuroblastoma phenotypes, including growth inhibition and differentiation, and that BMP4 mRNA and protein were induced by Wnt3a/Rspo2. In this study, we characterise the phenotypic effects of BMP4 on neuroblastoma cells, demonstrating convergent induction of MSX homeobox transcription factors by Wnt and BMP4 signalling and BMP4-induced growth suppression and differentiation. Immunohistochemical analysis of BMP4 expression in primary neuroblastomas confirms a striking absence of BMP4 in poorly differentiated tumours, in contrast to high expression in ganglion cells. These results are consistent with a tumour suppressive role for BMP4 in neuroblastoma. RNA sequencing following BMP4 treatment revealed induction of Notch signalling, verified by increases of Notch3 and Hes1 proteins. Together, our data demonstrate for the first time Wnt-BMP-Notch signalling crosstalk associated with growth suppression of neuroblastoma.

## INTRODUCTION

The canonical Wnt signalling pathway is a critical regulator of differentiation, proliferation, stemness, and determination of cell fate. Deregulation of Wnt signalling contributes to many cancers, resulting from both activating mutations of the proto-oncogene *CTNNB1* (encoding β-catenin), and loss of function mutations in negative regulators of the pathway, such as *APC* (encoding Adenomatous Polyposis Coli). These changes result in elevated nuclear β-catenin, which acts a co-activator for TCF/LEF transcription factors and a pro-growth/survival gene expression programme, exemplified by transcriptional activation of oncogenes such as *MYC* and *CCND1* (1,2). Wnt signalling in cancer can display extensive crosstalk with other morphogenetic pathways such as bone morphogenetic protein (BMP) and Notch signalling (3).

Given the developmental origins of the childhood cancer neuroblastoma, it was reasonable to examine whether deregulated Wnt signalling might be one of its central oncogenic drivers. Neuroblastoma is derived from the sympathoadrenal lineage of the neural crest (4,5), and neural crest cell (NCC) fates are contingent on tightly regulated orchestration of many signalling pathways prompting neural crest induction and specification, including Wnt/β-catenin (6,7) and BMP signalling (8). Additionally, a well-characterized differentiation block of NCCs, resulting in neuroblastoma, depends on *MYCN* amplification and over-expression (9), with MYCN transcriptionally repressing genes required for sympathetic nervous system differentiation (10,11). In other developmental contexts, *MYCN* is known to be a Wnt-induced gene (12), circumstantially supporting the possible oncogenicity of Wnt signalling in neuroblastoma.

Our previous work demonstrated high-levels of the Leucine Rich Repeat Containing G Protein-Coupled Receptor 5 (LGR5) mRNA and protein in undifferentiated neuroblastomas and neuroblastoma cell-lines. and that LGR5 was also an upstream regulator of Mitogen-Activated Protein Kinase (MAPK) signalling in neuroblastoma (13). LGR5 canonically functions as an R-Spondin receptor and increases Wnt/β-catenin signalling amplitude (14). However, we found that Wnt3a/Rspo2 treatment of neuroblastoma cell-lines did not lead to induction of *MYCN*, in fact MYCN and MYC protein levels decreased with Wnt3a/Rspo2 treatment, in contrast to previous reports suggesting induction of *MYC* in non-*MYCN* amplified (non-MNA) neuroblastomas due to Wnt/β-catenin signalling (15). Further phenotypic analysis of Wnt3a/Rspo2 treated neuroblastoma cell-lines revealed that Wnt/β-catenin signalling exerted context-dependent effects, including growth suppression and differentiation evident in SK-N-BE(2)-C and SH-SY5Y cells. In order to understand the Wnt-induced phenotypic changes, we conducted RNA sequencing and identified 90 high-confidence Wnt/β-catenin signalling target genes in SK-N-BE(2)-C cells. Bioinformatic analysis of these neuroblastoma Wnt target genes in primary tumour datasets showed that the 90 genes contained 4 distinct Wnt gene modules, or metagenes, the expression of which correlated with prognosis. Wnt metagenes 1 and 2, containing approximately 56% of our neuroblastoma Wnt target genes were expressed at markedly lower levels in high-risk neuroblastomas suggesting that these genes likely encode growth-suppressive and/or pro-differentiation proteins (16). Consistent with this idea, some genes with documented tumour-suppressive roles in neuroblastoma were included in these Wnt modules. These include *EPAS1* (17) and *MSX1* (18), both of which have been shown to inhibit neuroblastoma cell growth. Further analysis of our Wnt differentially expressed genes (DEGs) suggested that Wnt signalling is also a key regulator of mesenchymal and adrenergic differentiation states (19) which contribute to neuroblastoma cellular heterogeneity in both cell-lines and primary tumours (20,21).

One of our most highly Wnt-induced genes following Wnt induction was *BMP4*. Previous studies have alluded to a role for BMPs in neuroblastoma growth and differentiation, including BMP2 in mouse neuroblastoma and SH-SY5Y cells (22,23) and BMP4, which also affected SH-SY5Y differentiation and decreased proliferation markers (24). However, the mechanisms and signalling crosstalk involved in BMP-mediated phenotypes and the significance to primary disease of BMPs is not known. Given the interplay of Wnt and BMP signalling in neural crest development (25,26) and in many cancers (27-29), we hypothesised that the Wnt-BMP pathway may be key in regulating neuroblastoma growth. Specifically, although BMPs can have context-dependent roles in cancer, either promoting or inhibiting growth (30), Wnt-BMP signalling may be at the core of a growth-restrictive module in neuroblastoma, possibly via convergence on MSX transcription factors, which are known to be downstream of BMPs in neural crest specification (31) and also inhibit growth of neuroblastoma cells (18).

In this study we examine the relationship between Wnt and BMP signalling in neuroblastoma using functional assays and next-generation sequencing. Our study establishes co-ordinate signalling pathways operational in neuroblastoma which may be exploited for prognosis and therapeutics.

## MATERIALS AND METHODS

### Neuroblastoma cell lines and culture conditions

Neuroblastoma cell lines were a purchased from the European Collection of Authenticated Cell Cultures (ECACC) and from Deutsche Sammlung von Mikroorganismen und Zellkulturen (DSMZ). Their identity was verified by using STR profiling (Eurofins) and lack of Mycoplasma infection was confirmed by Mycoalert Mycoplasma Detection Kit (Lonza). Cell lines were cultured in Dulbecco’s modified eagle’s medium (DMEM):F12-HAM (Sigma) supplemented with 10% (v/v) foetal bovine serum (FBS) (Life technologies), 2mM L-glutamine, 100 U/mL penicillin, 0.1 mg/mL streptomycin, and 1% (v/v) non-essential amino acids. BMP4 (R&D Systems) treatment was carried out at low serum conditions (1-5 % FBS) in the same growth media.

### Incucyte live cell imaging and cell cycle analysis

Cell proliferation was monitored real time by using Incucyte Live Cell Imaging system using cell confluence as surrogate for growth, according to manufacturer’s instructions. Propidium-iodide labelling and fluorescence activated cell sorting (FACS) analysis to detect cell cycle phases was performed as previously described (32).

### Protein Extraction and Western blot

Cells were lysed in Radioimmunoprecipitation assay (RIPA) buffer and protein concentration was determined by using Micro BCA TM protein assay kit (Thermo Fisher). Western blot was performed as described previously (32). The antibodies used are listed in Supplementary Table 1.

### RNA extraction, reverse transcription and qPCR

RNA was extracted by using the miRNeasy kit (QIAGEN) and subsequently transcribed with Superscript IV (Invitrogen). Quantitative PCR was performed by using QuantiNova kit on Mx3500P (Stratagene). The oligonucleotide primers used in this study are shown in Supplementary Table 2.

### Immunohistochemistry

Tissue microarrays (TMAs), containing 47 pre-chemotherapy, peripheral neuroblastic tumours were stained by using a BMP4 antibody (EPR6211, Abcam). Immunohistochemistry staining was scored as positive or negative by a pathologist blinded to the specimens. All human tissues were acquired with appropriate local research ethics committee approval. Immunohistochemistry was performed with a Leica Microsystem Bond III automated machine using the Bond Polymer Refine Detection Kit (Ref DS9800) followed by Bond Dab Enhancer (AR9432). The slides were dewaxed with Bond Dewax Solution (AR9222). Heat mediated antigen retrieval was performed using Bond Epitope Retrieval Solution for 20 mins.

### RNA-seq and bioinformatic analysis

IMR32 cells were treated with 5 ng/mL BMP4 and 2% BSA (PBS) vehicle as control for 24 hours and were subsequently harvested. RNA was extracted by using miRNeasy Mini Kit (Qiagen), according to manufacturer’s instructions. Libraries were constructed and sequenced as previously described (16). Briefly, cDNA libraries were prepared from 1 ug RNA (TruSeq Stranded Total RNA Library Prep Kit, Illumina) and 100 bp, paired end reads were sequenced on Illumina HiSeq 2000. The reads were aligned to the human genome (hg38) by using STAR and the alignment files (BAM) files were further analysed in SeqMonk v1.45. (https://www.bioinformatics.babraham.ac.uk/projects/seqmonk/). Gene expression was quantified by using the Seqmonk RNA-seq analysis pipeline. Differentially expressed genes (DEG) were identified by DESEQ2 (p<0.005). sets, and a minimum fold difference threshold of 1.3 was applied. RNA sequencing data is available from the European Nucleotide Archive (ENA) under the study accession number PRJEB36530. We performed Gene Signature Enrichment Analyses (GSEAs) on preranked lists of log2-transformed relative gene expression values (Broad Institute). Kaplan Meier survival analysis and K means clustering were performed by using the R2 Genomics Analysis and Visualization Platform (http://r2.amc.nl).

## RESULTS

### Cross-talk between Wnt and BMP signalling in neuroblastoma

In our RNA-seq analysis of Wnt3a/Rspo2-treated neuroblastoma cells, *BMP4* was amongst the most highly induced genes (>50-fold) and BMP4 protein was also upregulated (16), leading us to further analyse global effects of Wnt3a/Rspo2-treatment on BMP/TGF genesets. As shown in Figure 1A, a volcano plot of Gene Set Enrichment Analysis (GSEA) highlights the activation of all BMP/TGF-related functional gene modules in comparison to the C2 collection of gene sets in the Molecular Signatures Database (MSigDB, Broad Institute). We verified several receptors and ligands included in these gene sets by qRT-PCR. Although BMP4 was the most highly induced ligand gene, *BMP2, BMP6* and *BMP7* also showed 2-3 fold induction and genes encoding receptors *BMPR1A* and *BMPR2* were upregulated by 30% (Figure 1B).

**Figure 1.**
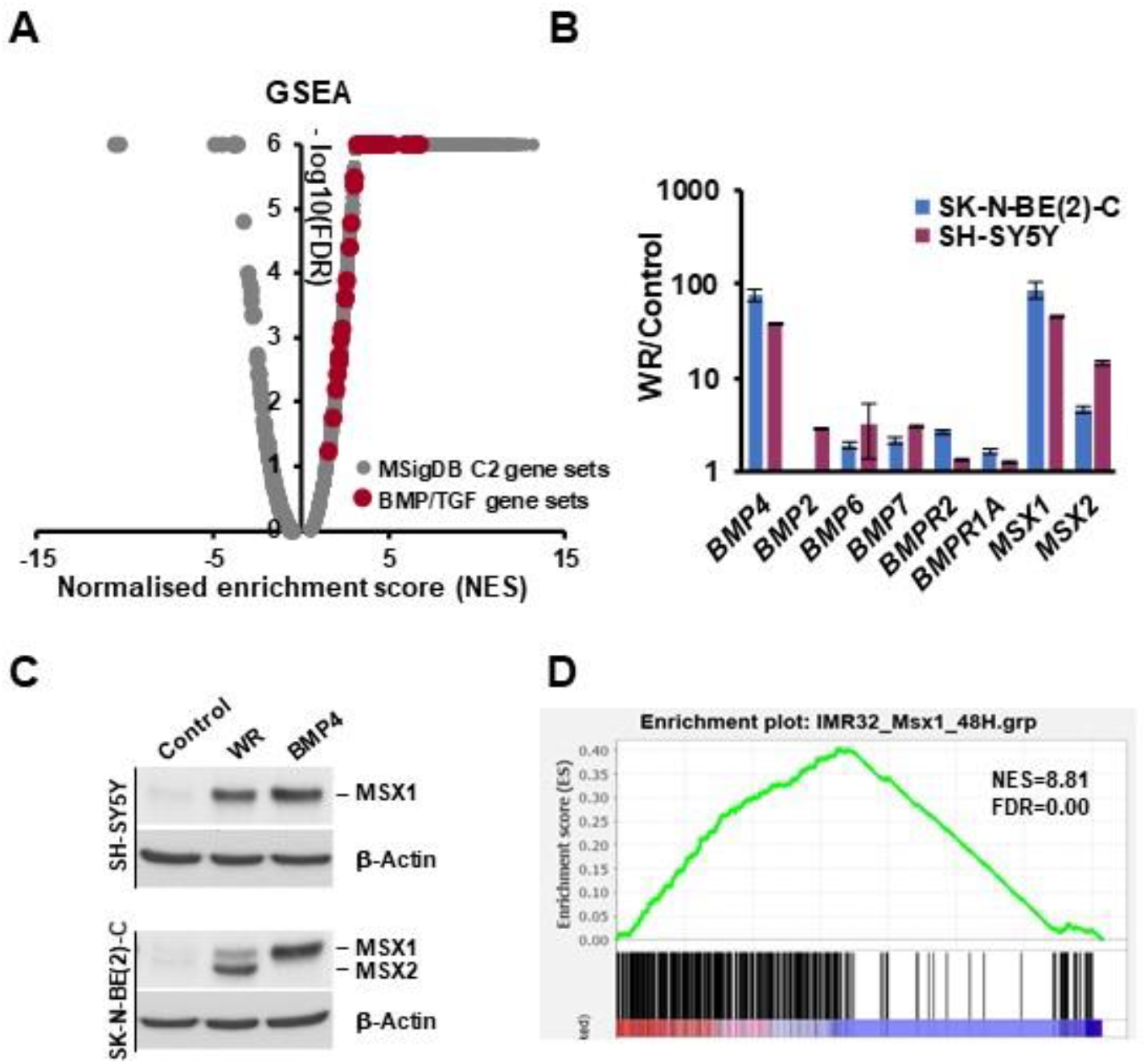
Convergence of Wnt and BMP pathways in NB. **(A)** GSEA Volcano plot on RNA-seq data of Wnt3a/RSPO-2 (WR) induced SK-N-BE(2)-C cells (ERP023744) with C2 group of gene sets (Molecular Signatures Database, Broad Institute), highlighting upregulation of BMP and TGF-β sets, shown in red. **(B)** Genes coding for BMP ligands, receptors and BMP target genes *MSX1/2*, are upregulated by 72H WR treatment in SK-N-BE(2)-C and SH-SY5Y cells. **(C)** Western blot of neural crest master regulators MSX1 and 2 induction by both WR and BMP4 treatment in neuroblastoma cells after 72-96H treatments. **(D)** GSEA showing upregulation of MSX1 target genes (GSE16481, 48H induction) in WR-treated SK-N-BE(2)-C.

*MSX1* and *MSX2*, which are known to be downstream of BMP4 (31), were also clearly induced, prompting us to assess the effects of Wnt3a/Rspo2 and BMP4 treatments at the protein level, using an antibody that recognises both MSX proteins. As seen in Figure 1C, analysis of treatments of 2 neuroblastoma cell lines consistently demonstrated that both Wnt and BMP4 ligands were able to strongly induce the MSX transcription factors, albeit to different degrees and with some selectivity of the paralogues induced apparent. Consistent with the convergence of Wnt and BMP signalling on MSX proteins, GSEA analysis confirmed a profound effect of Wnt3a/Rspo2 on the MSX1-regulated neuroblastoma transcriptome (Figure 1D).

Taken together, our data support Wnt and BMP4 signalling co-operating to regulate the neuroblastoma transcriptome, at least in part by convergence on MSX induction.

### BMP4 signalling affects growth and differentiation of neuroblastoma cells

We had previously demonstrated that Wnt ligands could inhibit the growth of neuroblastoma cells, and also influence their differentiation state (16,19). Given the intersection of Wnt and BMP signalling suggested by our transcriptomic data, we next sought to directly examine the phenotype of neuroblastoma cell-lines treated with BMP4. SK-N-BE(2)-C cells treated with as little as 0.1ng/ml BMP4 exhibited markedly decreased proliferation, with concentrations ranging between 1ng/ml – 50ng/ml essentially arresting cell-growth (Figure 2A-C). Activation of the BMP/TGF pathway was confirmed by the robust elevation of phospho-SMAD1/5/9. There was a marked increase of Tropomyosin receptor kinase A (TrkA), a well-established marker of good prognosis in neuroblastoma (33). Although immunoblotting with cleaved caspase 3 antibody showed that there was no apoptosis, an increase of the cell-cycle inhibitor p27 was evident, together with a decrease of MYCN and E2F1 proteins (Figure 2D). These markers are indicative of G1/S-phase cell cycle arrest, and this was confirmed by cell-cycle analysis (Supplementary Figure 1).

**Figure 2.**
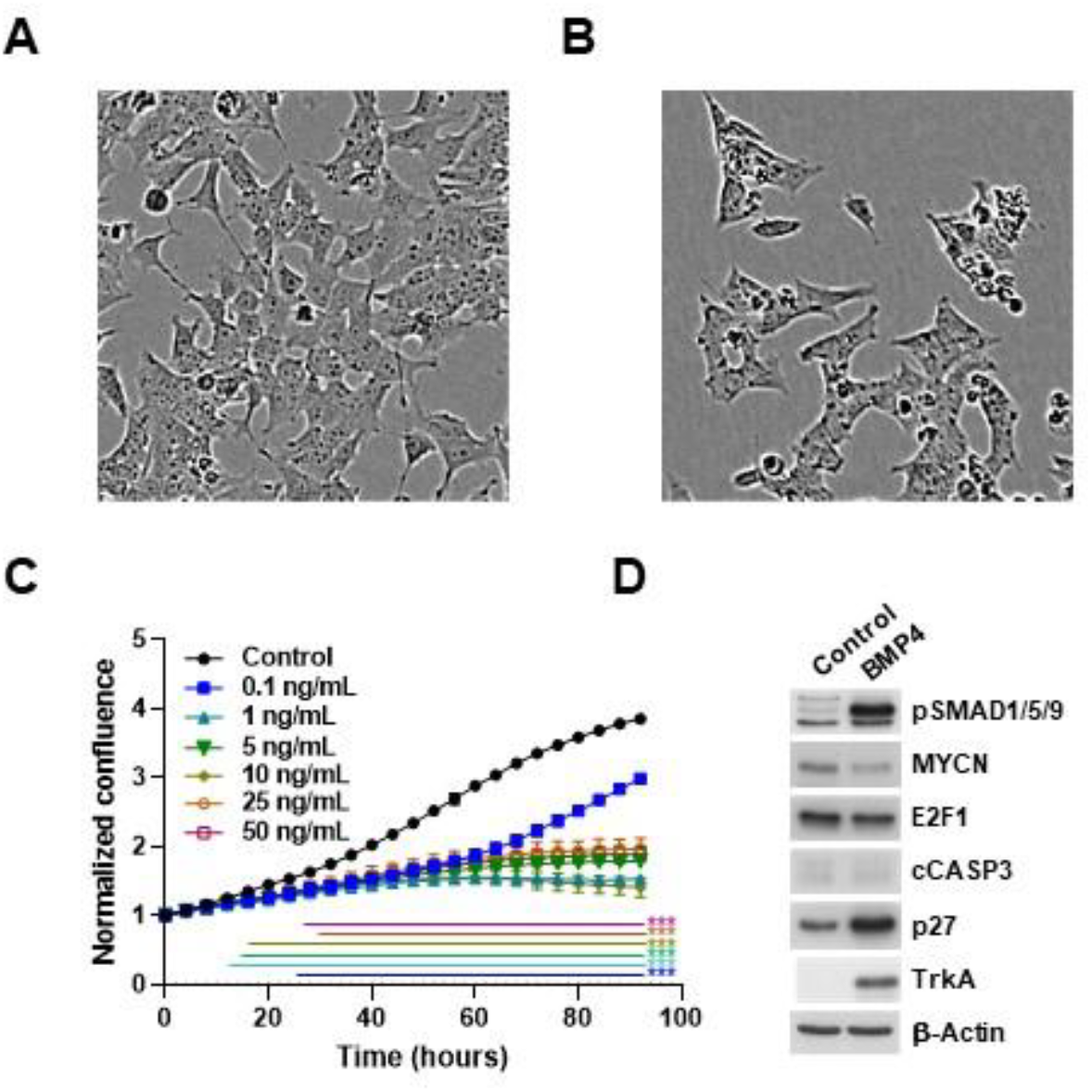
BMP4 treatment induces growth inhibition of SK-N-BE(2)-C cells. **(A)** Phase contrast image of vehicle-treated SK-N-BE(2)-C cells, and **(B)** 10 ng/mL BMP4-treated SK-N-BE(2)-C cells after 96 hours. **(C)** Cell confluence was measured and analysed in triplicates by Incucyte live cell imaging and was used as a surrogate for growth. Normalized confluence was significantly (p<0.01) reduced by BMP4 in all concentrations tested by 30 hours after treatment. Data is a representative of 2 biological replicates. **(D)** Western blot of changes in protein expression and phosphorylation after 96 hours of BMP4 treatment

We next assessed a second *MYCN*-amplified (MNA) neuroblastoma cell-line, IMR32, confirming the growth-inhibitory effects of BMP4, with the lowest significant effect observed at 5ng/ml (Figure 3). In general, growth suppression was not as marked in IMR32 cells relative to SK-N-BE(2)-C cells; however we observed clear neuritogenesis at concentrations of 1ng/ml and above (Figure 3B, D). Protein analysis again confirmed a robust phosphorylation of SMAD1/5/9, a lack of apoptosis, and induction of p27 and TrkA (Figure 3E). E2F1 was again decreased, but MYCN levels were not markedly affected. Consistent with the increase of neurites, dopamine β-hydroxylase (DBH) protein expression was induced by BMP4 treatment. In order to evaluate whether the effects of BMP4 were restricted to MNA neuroblastoma only, we also assessed the non-MNA neuroblastoma cell-line SH-SY5Y, and found that, similar to SK-N-BE(2)-C cells, BMP4 induced growth suppression. Like SK-N-BE(2)-C, SH-SY5Y cells showed no overt signs of neuritogenesis (Supplementary Figure 2). Four other neuroblastoma cell-lines (two MNA and two non-MNA) also showed similar growth suppression when treated with BMP4 (data not shown).

**Figure 3.**
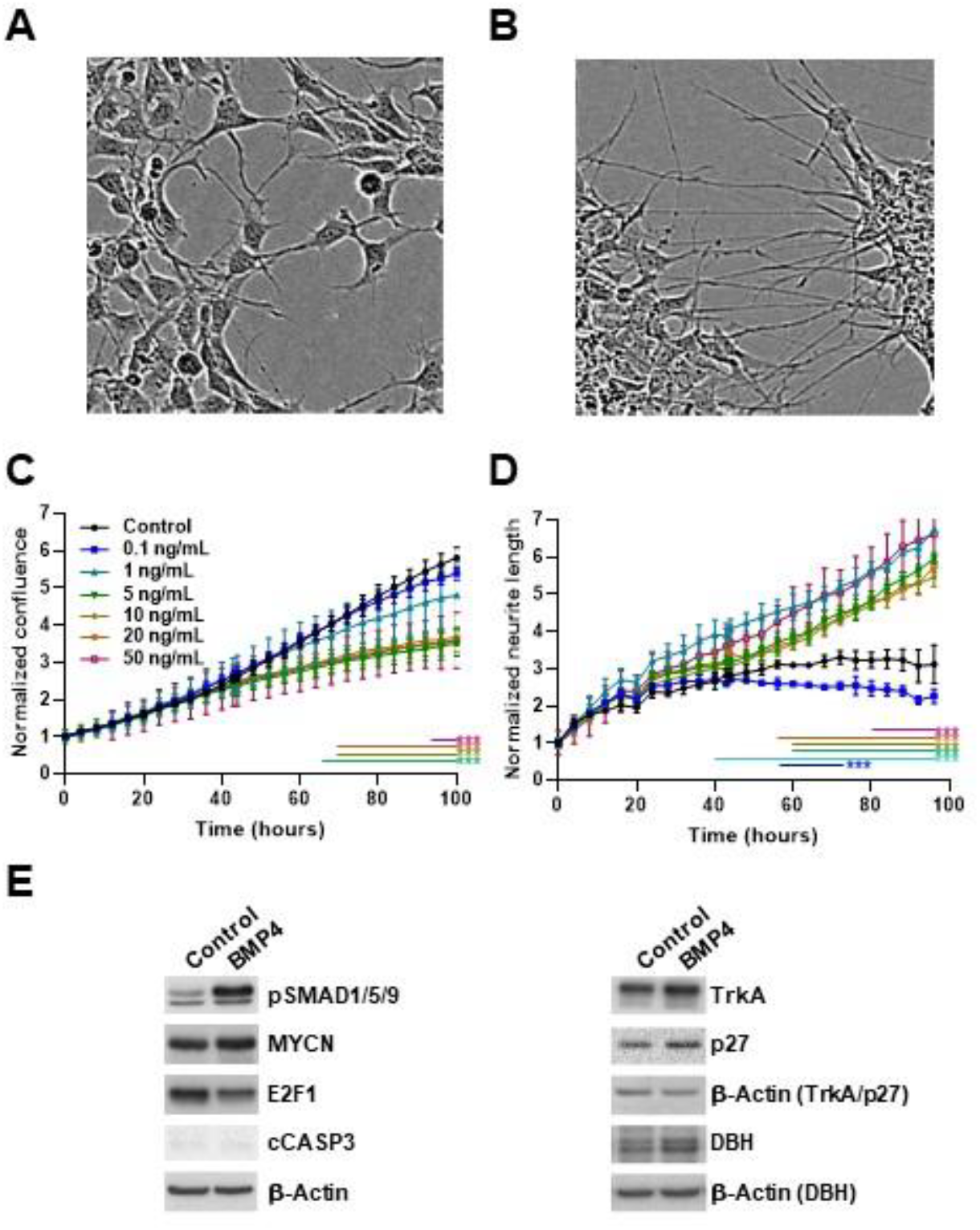
BMP4 treatment induces growth inhibition and neuritogenesis of IMR32 cells. Phase contrast image of **(A)** vehicle and **(B)** BMP4-treated IMR32 cells after 96 hours (10 ng/mL). **(C)** Normalized cell confluence, measured in triplicates, was significantly (p<0.01) reduced by BMP4 in concentrations above 5 ng/mL. Data is a representative of 3 biological replicates. **(D)** Significant changes (p<0.01) in normalized neurite length was observed after treatment with BMP4 at concentrations above 1 ng/mL. **(E)** Western blots showing protein expression and phosphorylation changes after 48-72 hours BMP4 treatment.

Together, these phenotypic analyses show that BMP signalling can block the proliferation of neuroblastoma cells, as well as inducing differentiation, similar to our findings with Wnt signalling in neuroblastoma (16).

### BMP4 protein expression correlates with better prognosis of neuroblastoma patients

It is well-known that BMP signalling can have both pro- and anti-proliferative effects, depending on cell context (30). Having established that BMP4 can restrict neuroblastoma proliferation and induce differentiation, we next sought to establish whether BMP4 expression in primary tumours would also reflect a potential tumour-suppressive function. Immunohistochemistry on neuroblastoma tissue-microarrays (TMAs) containing 47 neuroblastic tumour patient cores of different stages revealed that BMP4 expression was markedly restricted to tumours with more differentiation (ganglioneuroblastoma and ganglioneuroma), with virtually no expression of BMP4 evident in poorly differentiated neuroblastomas, including both *MYCN* amplified and non-amplified tumours (Figure 4A-C). Positive and negative controls for immunostaining are shown in Supplementary Figure 3. Statistical analysis confirmed the marked association of BMP expression with level of differentiation (p=0.0002, Figure 4D). Consistent with this, decreased *BMP4* mRNA in the transgenic *Th*-*MYCN*-driven mouse neuroblastoma model, relative to normal ganglia, was also revealed by analysis of a published dataset (Supplementary Figure 3D) (34). From the survival data available we could also ascertain a significant association (p=0.013) of low BMP4 expression and poor survival (Figure 4E). Thus the BMP4 expression pattern in primary tumours aligns with our *in vitro* functional analyses, strongly suggesting a pro-differentiation and anti-growth role for BMP signalling, particularly BMP4, in neuroblastoma.

**Figure 4.**
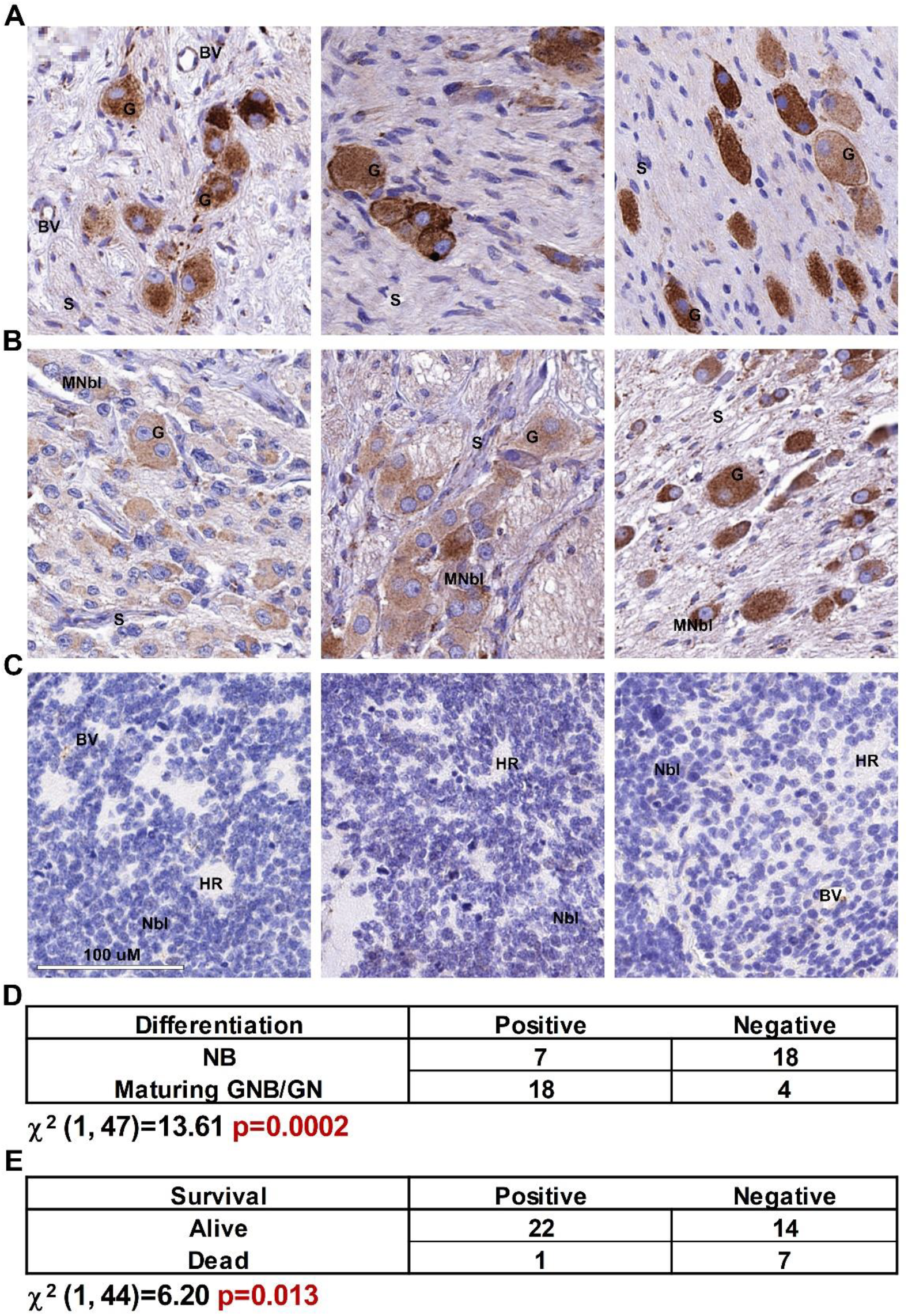
Immunohistochemical analysis of BMP4 protein expression in neuroblastic tumours. Expression of BMP4 protein in **(A)** mature ganglioneuroma, **(B)** ganglioneuroblastoma and maturing ganglioneuroma, **(C)** poorly differentiated neuroblastoma, where the third tumour is *MYCN*-amplified. BMP4 expression was positively and significantly correlated to **(D)** the degree of differentiation and **(E)** survival (Chi-squared test). Neuroblasts (Nbl), maturing neuroblasts (MNbl), ganglion cells (G), Schwannian stroma (S), Homer Wright rosettes (HR) and positively-staining blood vessels (BV) are indicated.

### Wnt and BMP4 signalling have overlapping but distinct effects on the neuroblastoma transcriptome

In order to better define the genes and pathways affected by Wnt and BMP pathways in neuroblastoma cells, we conducted RNA sequencing of IMR32 cells treated with BMP4. We identified 772 upregulated and 559 down-regulated genes (collectively referred to as BMP4 DEGs) after 24 hrs of BMP4 treatment. Known targets of BMP signalling such as *NOG, GREM2, GREM1* and *BAMBI* were strongly induced, together with the *ID* family of transcription factors, further verifying a functional, canonical BMP signalling pathway in IMR32 cells. Several Notch pathway genes were strongly induced, including *NOTCH3, HES1* and *HEY1* (Figure 5A). We found a highly significant overlap (p<0.001) of 31 genes between the 90 high confidence Wnt-regulated genes we identified previously in SK-N-BE(2)-C cells (16) and the 1331 BMP4-regulated genes (p=<0.005, DESEQ2 test, min. fold change 1.3), although 11 genes were oppositely regulated The complex interplay of BMP and Wnt signalling is also demonstrated by the strong but opposite regulation of the non-canonical Wnt ligand *WNT11* (Figure 5B).

**Figure 5.**
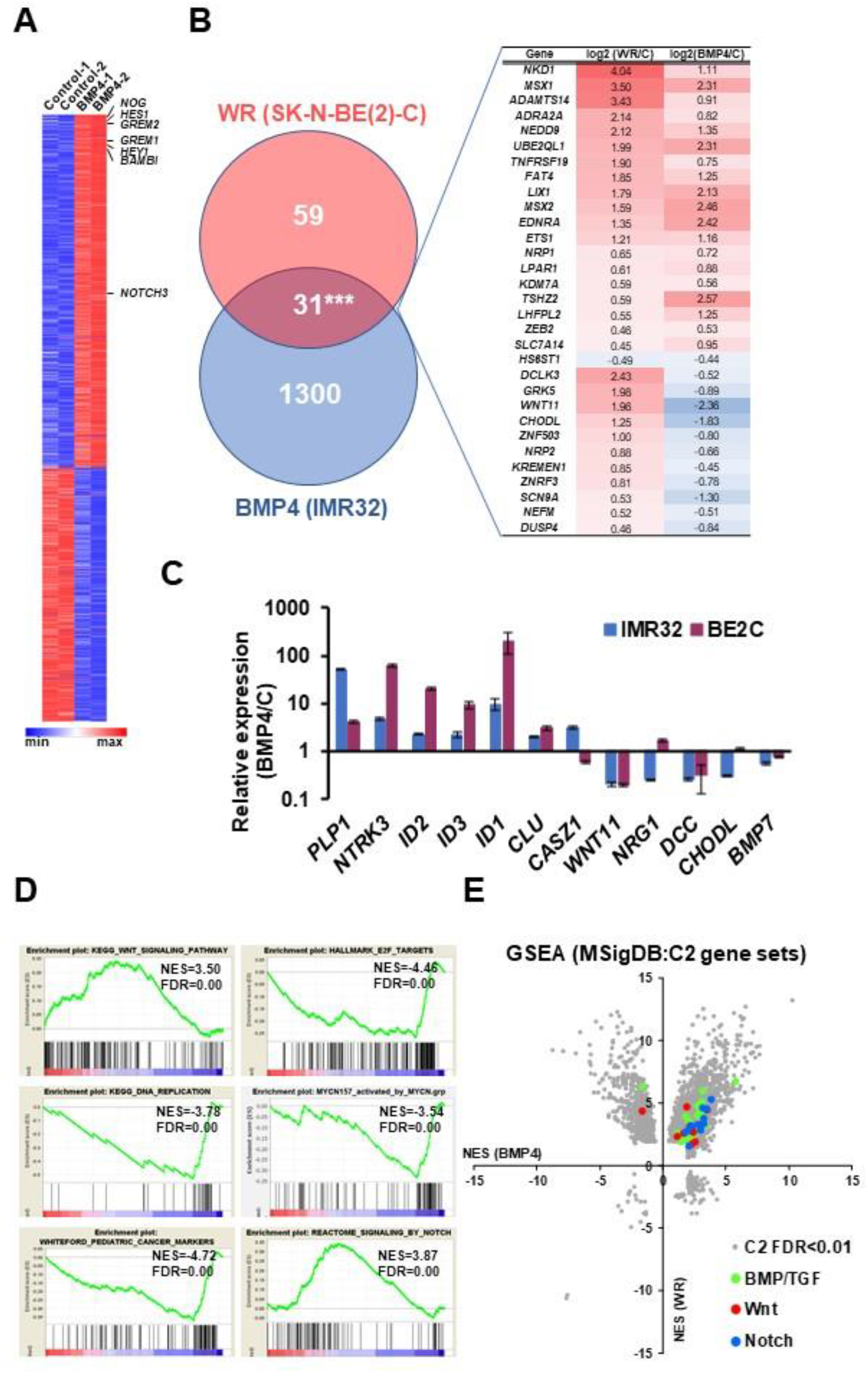
RNA-seq analysis of BMP4-treated IMR32 cells reveals Wnt and BMP4 signalling crosstalk in neuroblastoma cells. **(A)** Heatmap of DEGs in IMR32 after 24H BMP4 treatment (5 ng/mL), p<0.005 (DESEQ2), minimum fold change 1.3. Canonical BMP4 and Notch target genes are indicated. **(B)** Venn diagram and heatmap of shared target genes of BMP4 and WR datasets in neuroblastoma cells. The number of shared genes is significantly higher than expected by chance (hypergeometric test p=4.8e-22). **(C)** Validation of BMP4 target genes in 2 neuroblastoma cell lines. **(D)** GSEA highlighting significant regulation of functional gene sets. **(E)** Comparative GSEA analysis of C2 gene sets (Molecular Signatures Database, Broad Institute) in BMP4-treated IMR32 and WR-treated SK-N-BE(2)-C cells reveals upregulation of BMP/TGF (n=29), Wnt (n=12) and NOTCH (n=11) gene sets by both treatments.

In order to confirm that the IMR32 BMP4 regulated genes are representative of neuroblastoma generally, we validated a panel of 5 upregulated and 5 downregulated BMP4 targets in SK-N-BE(2)-C and IMR32 cells following BMP4 treatment. Strong concordance in BMP4 response was apparent between both cell lines (Figure 5C). Interestingly, the epigenetically silenced neuroblastoma tumour suppressor gene *CLU* was induced in both cell lines, together with another tumour suppressor *CASZ1* only in IMR32 cells (35,36). GSEA revealed that BMP4 induced strong upregulation of Wnt pathway genes in IMR32, as well as downregulation of E2F1 and DNA replication signatures (Figure 5D). We also constructed a gene set based on a functional neuroblastoma-specific MYCN signature of 157 genes whose up- or down-regulation in IMR32 cells had been demonstrated to be strongly linked with neuroblastoma prognosis (MYCN157) (37). BMP4 treatment had a repressive effect on the MYCN-activated members of the MYCN157 signature and also negatively regulated pediatric cancer markers. Upregulation of the Notch signalling pathway genes was confirmed by GSEA (Figure 5D). In order to examine a potential crosstalk of signalling pathways at a global level, we plotted normalized enrichment scores (NES) of GSEAs for our previous Wnt RNA-seq together with our BMP4 RNA-seq data, using the C2 collection of Molecular Signatures Database (Broad Institute). We found not only a strong mutual upregulation of genes participating in Wnt and BMP signalling, but also Notch signalling with both BMP and Wnt (Figure 5E).

We had previously utilised our Wnt DEGs for k-means clustering of RNA-seq expression data of primary neuroblastoma tumours (SEQC, GSE62564 (38)) to demonstrate that they are capable of segregating clinical subtypes of neuroblastoma, such that a Wnt DEGs-dependent low risk (Wnt-LR), intermediate risk (Wnt-IR), and two high-risk (with or without MNA, Wnt-HR) categories could be identified (16). We therefore tested whether the expression patterns of our BMP4 DEGs, which included both up and downregulated genes, in primary tumours might also be able to partition clinical subtypes and be predictive of outcome. As shown in Figure 6A, k-means clustering of RNA-seq data of 498 primary neuroblastomas (SEQC) with the 1331 BMP4 DEGs identified 3 patient clusters, corresponding to 3 distinct risk categories. Kaplan-Meier curves for the three BMP clusters demonstrated that BMP4 cluster 1 was associated with low-risk, BMP4 cluster 2 with intermediate risk, and BMP4 cluster 3 with high risk (Figure 6B). Further, BMP4 cluster 1 aligned remarkably closely with Wnt-LR, BMP4 cluster 2 with Wnt-IR, and BMP4 cluster 3 with Wnt-HR (p=1.1e-150 Chi-squared test, Supplementary Figure 4). Kaplan-Meier analysis of BMP4 upregulated genes showed that their high expression correlates with good prognosis, whereas high expression of BMP4 downregulated genes correlates with poor prognosis (Figure 6C-D), suggesting that BMP4 induces gene expression programmes associated with less aggressive clinical subtypes.

**Figure 6.**
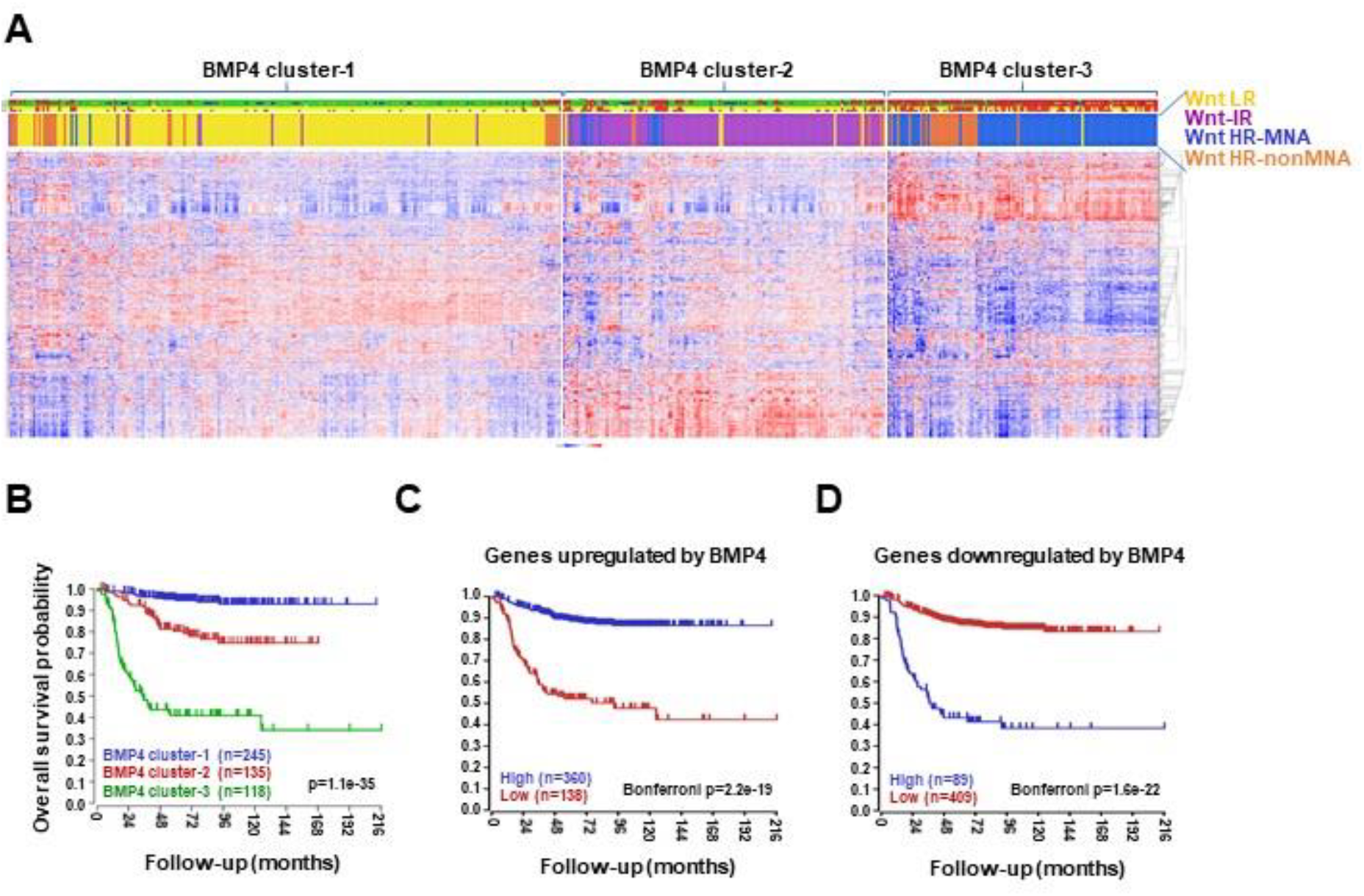
Meta-analysis of BMP4 target gene expression in primary neuroblastomas. **(A)** K-means clustering of BMP4 target genes in primary tumour gene expression data set SEQC (n=498, k=3). Coloured bars on the top indicate risk status, stage, MYCN amplification, survival, progression and clustering according to WR target genes. **(B)** Clustering according to BMP4 target gene expression divided the SEQC patient cohort into prognostic groups with significantly different survival probabilities. **(C)** Kaplan-Meier analyses showing that high expression of BMP4-upregulated genes strongly and significantly correlates with survival, while **(D)** high levels of genes downregulated by BMP4 is associated with poor prognosis.

Taken together, our RNA-seq reveals profound effects of BMP4 on the neuroblastoma cell transcriptome, including interaction with Wnt and Notch signalling. Importantly, our analyses strongly support a growth-suppressive function for BMP4 signalling in neuroblastoma.

### Notch signalling is downstream of Wnt-BMP signalling

Our analysis of Wnt DEGs had suggested interactions between Wnt and Notch signalling in neuroblastoma (16). As MSX1 has been shown to upregulate Notch signalling (18), we next examined the links between Wnt, BMP4, MSX1 and Notch in neuroblastoma. Notch pathway targets and effectors were highly induced in our RNA-seq of IMR32 treated with BMP4, so we first validated a panel of Notch pathway genes, including *HES1, MAFB, MAML2* and *NOTCH3* (Figure 7A), confirming a BMP4-Notch signalling pathway in neuroblastoma. We further examined BMP4 effects on Notch signalling by protein analysis of IMR32 and SK-N-BE(2)-C cell lysates. In both cell-lines, simultaneous induction of MSX1 and cleavage of NOTCH3 was observed, together with upregulation of the Notch target and effector protein HES1 (Figure 7B). Real-time PCR confirmed that BMP4 induced Notch genes in SK-N-BE(2)-C cells too (Figure 7C). Finally, we evaluated the regulation of NOTCH3-ICD induced genes identified in IMR32 cells (39) in our BMP4 and Wnt RNA-seq data. GSEA revealed a striking positive correlation between BMP4 and NOTCH-ICD induced genes, and similarly with BMP4 and NOTCH-ICD repressed genes (Figure 7D). Our Wnt DEGs also showed a correlation with the NOTCH3-ICD regulated transcriptome in neuroblastoma, but to a lesser extent than BMP4 (Figure 7E). Together these studies confirm the strong interplay of Wnt, BMP and Notch signalling in determining transcriptional programmes in neuroblastoma, leading to growth arrest and, in some cell-lines, differentiation (Figure 8).

**Figure 7.**
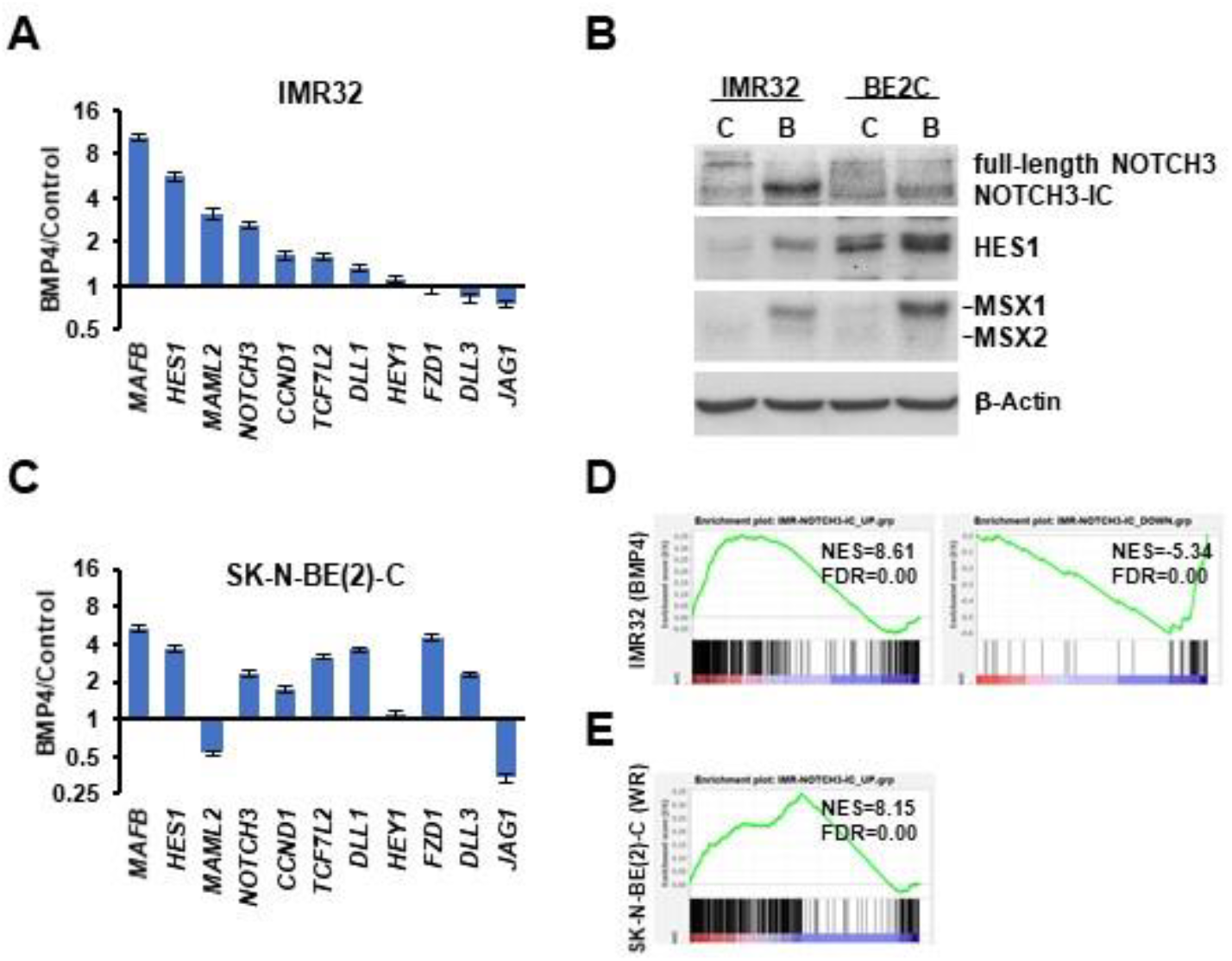
BMP4 and Wnt regulate Notch signalling in neuroblastoma. **(A)** Regulation of Notch pathway genes by BMP4 in IMR32 cells, detected by qPCR. **(B)** BMP4 treatment leads to NOTCH3 cleavage, upregulation of Notch target/effector protein HES1 and MSX1/2 proteins in IMR32 and SK-N-BE(2)-C cells. **(C)** Regulation of Notch pathway genes by BMP4 in SK-N-BE(2)-C cells, detected by qPCR. **(D)** GSEA of RNA-seq data sets of BMP4-treated IMR32 and **(E)** WR-induced SK-N-BE(2)-C with NOTCH3-IC target gene sets identified in IMR32.

**Figure 8.**
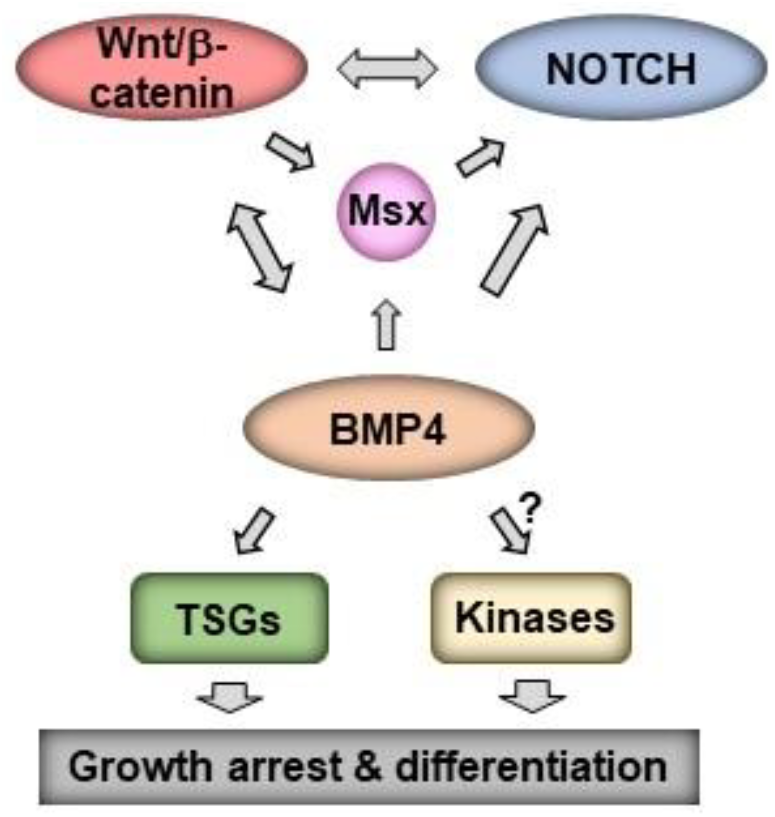
Model for Wnt, BMP, MSX and Notch crosstalk. Regulatory interactions promoting growth arrest and differentiation in neuroblastoma are outlined.

## DISCUSSION

The molecular etiology of neuroblastoma has been intensively studied at the level of deregulated transcriptomes resulting from altered developmental transcription factor expression, as exemplified by MYCN. Perhaps surprisingly for an embryonic tumour associated with a disrupted differentiation programme, our understanding of the signalling pathway networks that are involved in neuroblastoma tumorigenesis remains relatively limited. In particular, the role of the Wnt signalling pathway, often clearly oncogenic in both adult and childhood cancers such as colorectal cancer and Wilms’ tumour (1), has been found to exert more complex influences in NB, as shown several laboratories, including ours. These data have indicated Wnt-induced growth promotion or suppression (13,16,40,41), modulation of signalling and transcriptional pathways, interactions with MYCN (16,41), and underlying changes in differentiation states contributing to neuroblastoma tumour heterogeneity (16,19). In this study, we define a Wnt-BMP growth inhibitory axis in neuroblastoma, identifying MSX and Notch signalling as downstream mediators of growth suppression.

Previous studies have suggested a role for BMP signalling in regulating neuroblastoma cell differentiation and proliferation. However, the upstream and downstream mechanistic intermediates of BMPs have never been characterized, including the crosstalk with other signalling pathways. BMP2 induced neural differentiation in mouse neuroblastoma cells by increasing the expression of neurogenic transcription factors *Dlx2, Brn3a*, and *NeuroD6* (22). Similarly, BMP2 was shown to induce growth arrest and neuronal differentiation of SH-SY5Y and RTBM cell-lines subsequent to accumulation of p27 (23). Although *BMP2* has been shown to be a Wnt target gene in other cell types, our RNA-seq of Wnt3a/Rspo2 treated neuroblastoma cells highlighted *BMP4* as the most prominent BMP gene downstream of Wnt/β-catenin signalling, and a recent study showed that BMP4 can trigger neurite-like extensions in SH-SY5Y cells, induce TrkA, and decrease MYCN and Ki-67 (24). Our study in part agrees with these findings, but with important differences and clearer mechanistic insights. Our studies quantify and confirm growth suppression of not only SH-SY5Y cells, but also SK-N-BE(2)-C and IMR32 cells, together with increases of TrkA. Clear neuritogenesis was only observed in IMR32 cells. Furthermore, our studies do not suggest that reduced MYCN levels are the predominant effect of BMP4 treatment, Ferlemann *et al* having suggested this based on MYCN reduction in the non-MNA neuroblastoma line SH-SY5Y (24). Given our confirmation of G1-phase cell-cycle arrest, reductions in levels of MYCN and E2F1 may be attributable to cell-cycle effects. However, based on GSEA analysis of our RNA-seq data, which demonstrate a strong effect of BMP4 on MYCN-mediated gene regulation, we suggest that BMP4 might more subtly interfere with MYCN transcriptional activity.

Our transcriptomic analysis did not support activation of *DLX2, BRN3A*, or *NEUROD6* genes as being involved in the BMP4-induced phenotypes of neuroblastoma cells, as previously suggested by studies using BMP2 (22). Rather, our studies confirmed strong and consistent induction of MSX proteins, in particular MSX1. MSX1 is already known to be downstream of BMP signalling in the developing neural crest (31) and has been shown to suppress proliferation and colony formation in soft agar when exogenously over-expressed in neuroblastoma cells. This study by Revet et al further established that MSX1 activated NOTCH3 and that low mRNA expression of the Notch pathway target *HEY1* correlated with poor prognosis (18). The suggested link between BMP, MSX and Notch signalling was verified by both our RNA-seq and protein expression analyses. The role(s) of Notch signalling in neuroblastoma are complex, and although recent studies have linked it primarily with tumour heterogeneity and differentiation states of neuroblastoma (21,42), other studies, in addition to Revet et al (18), have suggested that Notch signalling can be growth suppressive in neuroblastoma. In particular, neuroblastoma growth inhibition was demonstrated following exogenous expression of the intracellular domain of all three NOTCH proteins (NOTCH1-3). Furthermore, transduction with HES1 of several neuroblastoma cell-lines, including IMR32 and SH-SY5Y, led to inhibition of proliferation, and growth was also inhibited by treatment with recombinant Notch ligand Jag1 (43). Another study associating Notch signalling with cell migration reported that the NOTCH3-ICD led to a transient attenuation of the cell-cycle (39). Importantly, a study demonstrating the therapeutic benefits of the histone deacetylase inhibitor panobinostat *in vivo*, using the *Th*-*MYCN* mouse model of neuroblastoma, demonstrated that growth inhibition and differentiation resulting from panobinostat were accompanied by increased Notch and BMP pathway genes (44). On the basis of published work and our data, we propose that the net effect of BMP signalling in neuroblastoma is contingent on signalling crosstalk with Notch.

Whilst our analyses provide, for the first time, strong evidence for the interplay of Wnt, BMP4 and Notch signalling in neuroblastoma, it is unlikely that the profound effects of BMP4 on the neuroblastoma transcriptome and phenotype are mediated solely by any factor, but are rather a co-ordinated effect of the complex interactions of signalling pathways and developmental transcription factors. BMP4 can also control growth by non-SMAD-dependent pathways such as MEK/ERK signalling (45) and it will be important to conduct phosphoproteomic analyses in the future. Nevertheless, our compelling demonstration of BMP4 protein absence in poorly differentiated and aggressive neuroblastomas, and BMP4’s strong anti-proliferative effect raise the possibility of exploiting BMP4 or agonists of BMP signalling as therapeutic agents. Interestingly, a BMP9-derived peptide has been shown to enhance differentiation of neuroblastoma cells (46). Our characterization of the Wnt-BMP-MSX-Notch pathway and its associated biomarkers rationalises the evaluation of such novel therapeutics, as well as providing a foundation for delineating signalling regulatory networks in neuroblastoma.

**Supplementary figure S1. Cell-cycle analyses of BMP4 treated IMR32 and SK-N-BE(2)-C cells.** Propidium Iodide staining and flow cytometry reveal significant G1-arrest in IMR32 and SK-N-BE(2)-C cells following BMP4 treatment. (* p<0.05, *** p<0.01). Data for IMR32 represents two biological replicates.

**Supplementary figure S2. BMP4 treatment induces growth inhibition of SH-SY5Y cells.** Phase contrast image of **(A)** control and **(B)** BMP4-treated SH-SY5Y cells after 96 hours (10 ng/mL). **(C)** Normalized cell confluence was significantly (p<0.01) reduced by BMP4 treatment in concentrations above 1 ng/mL. **(D)** Western blots of protein expression, phosphorylation and cleavage after BMP4 treatment (10 ng/mL).

**Supplementary figure S3. BMP4 protein expression tissue controls for immunohistochemistry, and *Bmp4* expression in the *Th*-*MYCN* mouse neuroblastoma model. (A)** Colon (positive control), **(B)** Kidney (positive control), **(C)** Fat (negative control). **(D)** Boxplots showing decreased *Bmp4* mRNA in *Th-MYCN* neuroblastomas (GSE17740).

**Supplementary figure S4. Correlations between BMP4 and Wnt DEG prognostic clusters in 498 neuroblastomas.** Strong and significant correlation was observed between clustering of SEQC tumours according to BMP4 and Wnt target genes (Chi-squared test, p=1.1e-150).

## Abbreviations

BMP: Bone morphogenetic protein
EMT: epithelial-mesenchymal transition
ERK: extracellular signal–regulated kinase
GRN: gene regulatory network
GSEA: Geneset Enrichment Analysis.
ICD: Intracellular Domain.
KEGG: Kyoto encyclopedia of genes and genomes
LEF: lymphoid enhancer binding factor
MEK: MAPK/ERK kinase
MNA: *MYCN*-amplified
Rspo: R-spondin
TCF: T-cell-factor
TGF: Transforming Growth Factor

## ACKNOWLEDGEMENTS

We wish to thank Jane Coghill and Christy Waterfall at the University of Bristol Genomics Facility for help with transcriptomics, and Dr. Andy Herman for help with flow cytometry. We greatly indebted to Ms Aysen Yuksei and Dr Michael Krivanek for construction of the tissue microarrays and reviewing pathology data. We also thank the Children’s Cancer and Leukaemia Group (CCLG), the Biotechnology and Biological Sciences Research Council (BB/P008232/1), Cancer Research UK (A12743/A21046) and the Showering Fund for funding this study.

## AUTHOR CONTRIBUTIONS

**Marianna Szemes:** Conceptualization, Methodology, Formal analysis, Investigation, Data curation, Writing - original draft; review & editing. **Zsombor Melegh:** Formal analysis, Investigation. **Jacob Bellamy & Madhu Kollareddy:** Investigation. **Daniel Catchpoole:** Resources. **Karim Malik:** Conceptualization, Methodology, Formal analysis, Investigation, Funding acquisition, Supervision, Writing - original draft; review & editing.

## REFERENCES

1. Clevers, H. and Nusse, R. (2012) Wnt/beta-catenin signaling and disease. Cell, 149, 1192–1205.

2. Nusse, R. and Clevers, H. (2017) Wnt/beta-Catenin Signaling, Disease, and Emerging Therapeutic Modalities. Cell, 169, 985–999.

3. Katoh, M. (2007) Networking of WNT, FGF, Notch, BMP, and Hedgehog signaling pathways during carcinogenesis. Stem Cell Rev, 3, 30–38.

4. Brodeur, G.M. (2003) Neuroblastoma: biological insights into a clinical enigma. Nat Rev Cancer, 3, 203–216.

5. Johnsen, J.I., Dyberg, C. and Wickstrom, M. (2019) Neuroblastoma-A Neural Crest Derived Embryonal Malignancy. Front Mol Neurosci, 12, 9.

6. Garcia-Castro, M.I., Marcelle, C. and Bronner-Fraser, M. (2002) Ectodermal Wnt function as a neural crest inducer. Science, 297, 848–851.

7. Leung, A.W., Murdoch, B., Salem, A.F., Prasad, M.S., Gomez, G.A. and Garcia-Castro, M.I. (2016) WNT/beta-catenin signaling mediates human neural crest induction via a pre-neural border intermediate. Development, 143, 398–410.

8. Steventon, B., Araya, C., Linker, C., Kuriyama, S. and Mayor, R. (2009) Differential requirements of BMP and Wnt signalling during gastrulation and neurulation define two steps in neural crest induction. Development, 136, 771–779.

9. Brodeur, G.M., Seeger, R.C., Schwab, M., Varmus, H.E. and Bishop, J.M. (1984) Amplification of N-myc in untreated human neuroblastomas correlates with advanced disease stage. Science, 224, 1121–1124.

10. Westermark, U.K., Wilhelm, M., Frenzel, A. and Henriksson, M.A. (2011) The MYCN oncogene and differentiation in neuroblastoma. Seminars in Cancer Biology, 21, 256–266.

11. Gherardi, S., Valli, E., Erriquez, D. and Perini, G. (2013) MYCN-mediated transcriptional repression in neuroblastoma: the other side of the coin. Frontiers in oncology, 3, 42.

12. ten Berge, D., Brugmann, S.A., Helms, J.A. and Nusse, R. (2008) Wnt and FGF signals interact to coordinate growth with cell fate specification during limb development. Development, 135, 3247–3257.

13. Vieira, G.C., Chockalingam, S., Melegh, Z., Greenhough, A., Malik, S., Szemes, M., Park, J.H., Kaidi, A., Zhou, L., Catchpoole, D. et al. (2015) LGR5 regulates pro-survival MEK/ERK and proliferative Wnt/beta-catenin signalling in neuroblastoma. Oncotarget, 6, 40053–40067.

14. de Lau, W., Peng, W.C., Gros, P. and Clevers, H. (2014) The R-spondin/Lgr5/Rnf43 module: regulator of Wnt signal strength. Genes Dev, 28, 305–316.

15. Liu, X., Mazanek, P., Dam, V., Wang, Q., Zhao, H., Guo, R., Jagannathan, J., Cnaan, A., Maris, J.M. and Hogarty, M.D. (2008) Deregulated Wnt/beta-catenin program in high-risk neuroblastomas without MYCN amplification. Oncogene, 27, 1478–1488.

16. Szemes, M., Greenhough, A., Melegh, Z., Malik, S., Yuksel, A., Catchpoole, D., Gallacher, K., Kollareddy, M., Park, J.H. and Malik, K. (2018) Wnt Signalling Drives Context-Dependent Differentiation or Proliferation in Neuroblastoma. Neoplasia, 20, 335–350.

17. Westerlund, I., Shi, Y., Toskas, K., Fell, S.M., Li, S., Surova, O., Sodersten, E., Kogner, P., Nyman, U., Schlisio, S. et al. (2017) Combined epigenetic and differentiation-based treatment inhibits neuroblastoma tumor growth and links HIF2alpha to tumor suppression. Proc Natl Acad Sci U S A, 114, E6137–E6146.

18. Revet, I., Huizenga, G., Chan, A., Koster, J., Volckmann, R., van Sluis, P., Ora, I., Versteeg, R. and Geerts, D. (2008) The MSX1 homeobox transcription factor is a downstream target of PHOX2B and activates the Delta-Notch pathway in neuroblastoma. Exp Cell Res, 314, 707–719.

19. Szemes, M., Greenhough, A. and Malik, K. (2019) Wnt Signaling Is a Major Determinant of Neuroblastoma Cell Lineages. Front Mol Neurosci, 12, 90.

20. Boeva, V., Louis-Brennetot, C., Peltier, A., Durand, S., Pierre-Eugene, C., Raynal, V., Etchevers, H.C., Thomas, S., Lermine, A., Daudigeos-Dubus, E. et al. (2017) Heterogeneity of neuroblastoma cell identity defined by transcriptional circuitries. Nat Genet, 49, 1408–1413.

21. van Groningen, T., Koster, J., Valentijn, L.J., Zwijnenburg, D.A., Akogul, N., Hasselt, N.E., Broekmans, M., Haneveld, F., Nowakowska, N.E., Bras, J. et al. (2017) Neuroblastoma is composed of two super-enhancer-associated differentiation states. Nat Genet, 49, 1261–1266.

22. Du, Y. and Yip, H. (2010) Effects of bone morphogenetic protein 2 on Id expression and neuroblastoma cell differentiation. Differentiation, 79, 84–92.

23. Nakamura, Y., Ozaki, T., Koseki, H., Nakagawara, A. and Sakiyama, S. (2003) Accumulation of p27 KIP1 is associated with BMP2-induced growth arrest and neuronal differentiation of human neuroblastoma-derived cell lines. Biochem Biophys Res Commun, 307, 206–213.

24. Ferlemann, F.C., Menon, V., Condurat, A.L., Rossler, J. and Pruszak, J. (2017) Surface marker profiling of SH-SY5Y cells enables small molecule screens identifying BMP4 as a modulator of neuroblastoma differentiation. Sci Rep, 7, 13612.

25. Bhatt, S., Diaz, R. and Trainor, P.A. (2013) Signals and switches in Mammalian neural crest cell differentiation. Cold Spring Harb Perspect Biol, 5.

26. Martik, M.L. and Bronner, M.E. (2017) Regulatory Logic Underlying Diversification of the Neural Crest. Trends Genet, 33, 715–727.

27. Guo, X. and Wang, X.F. (2009) Signaling cross-talk between TGF-beta/BMP and other pathways. Cell Res, 19, 71–88.

28. Itasaki, N. and Hoppler, S. (2010) Crosstalk between Wnt and bone morphogenic protein signaling: a turbulent relationship. Dev Dyn, 239, 16–33.

29. Katoh, M. (2011) Network of WNT and other regulatory signaling cascades in pluripotent stem cells and cancer stem cells. Curr Pharm Biotechnol, 12, 160–170.

30. Bach, D.H., Park, H.J. and Lee, S.K. (2018) The Dual Role of Bone Morphogenetic Proteins in Cancer. Mol Ther Oncolytics, 8, 1–13.

31. Tribulo, C., Aybar, M.J., Nguyen, V.H., Mullins, M.C. and Mayor, R. (2003) Regulation of Msx genes by a Bmp gradient is essential for neural crest specification. Development, 130, 6441–6452.

32. Park, J.H., Szemes, M., Vieira, G.C., Melegh, Z., Malik, S., Heesom, K.J., Von Wallwitz-Freitas, L., Greenhough, A., Brown, K.W., Zheng, Y.G. et al. (2015) Protein arginine methyltransferase 5 is a key regulator of the MYCN oncoprotein in neuroblastoma cells. Molecular oncology, 9, 617–627.

33. Kramer, K., Gerald, W., LeSauteur, L., Uri Saragovi, H. and Cheung, N.K. (1996) Prognostic value of TrkA protein detection by monoclonal antibody 5C3 in neuroblastoma. Clin Cancer Res, 2, 1361–1367.

34. Balamuth, N.J., Wood, A., Wang, Q., Jagannathan, J., Mayes, P., Zhang, Z., Chen, Z., Rappaport, E., Courtright, J., Pawel, B. et al. (2010) Serial transcriptome analysis and cross-species integration identifies centromere-associated protein E as a novel neuroblastoma target. Cancer research, 70, 2749–2758.

35. Corvetta, D., Chayka, O., Gherardi, S., D’Acunto, C.W., Cantilena, S., Valli, E., Piotrowska, I., Perini, G. and Sala, A. (2013) Physical interaction between MYCN oncogene and polycomb repressive complex 2 (PRC2) in neuroblastoma: functional and therapeutic implications. J Biol Chem, 288, 8332–8341.

36. Wang, C., Liu, Z., Woo, C.W., Li, Z., Wang, L., Wei, J.S., Marquez, V.E., Bates, S.E., Jin, Q., Khan, J. et al. (2012) EZH2 Mediates epigenetic silencing of neuroblastoma suppressor genes CASZ1, CLU, RUNX3, and NGFR. Cancer research, 72, 315–324.

37. Valentijn, L.J., Koster, J., Haneveld, F., Aissa, R.A., van Sluis, P., Broekmans, M.E., Molenaar, J.J., van Nes, J. and Versteeg, R. (2012) Functional MYCN signature predicts outcome of neuroblastoma irrespective of MYCN amplification. Proc Natl Acad Sci U S A, 109, 19190–19195.

38. Zhang, W., Yu, Y., Hertwig, F., Thierry-Mieg, J., Zhang, W., Thierry-Mieg, D., Wang, J., Furlanello, C., Devanarayan, V., Cheng, J. et al. (2015) Comparison of RNA-seq and microarray-based models for clinical endpoint prediction. Genome biology, 16, 133.

39. van Nes, J., Chan, A., van Groningen, T., van Sluis, P., Koster, J. and Versteeg, R. (2013) A NOTCH3 transcriptional module induces cell motility in neuroblastoma. Clin Cancer Res, 19, 3485–3494.

40. Zins, K., Schafer, R., Paulus, P., Dobler, S., Fakhari, N., Sioud, M., Aharinejad, S. and Abraham, D. (2016) Frizzled2 signaling regulates growth of high-risk neuroblastomas by interfering with beta-catenin-dependent and beta-catenin-independent signaling pathways. Oncotarget, 7, 46187–46202.

41. Duffy, D.J., Krstic, A., Schwarzl, T., Halasz, M., Iljin, K., Fey, D., Haley, B., Whilde, J., Haapa-Paananen, S., Fey, V. et al. (2016) Wnt signalling is a bi-directional vulnerability of cancer cells. Oncotarget, 7, 60310–60331.

42. van Groningen, T., Akogul, N., Westerhout, E.M., Chan, A., Hasselt, N.E., Zwijnenburg, D.A., Broekmans, M., Stroeken, P., Haneveld, F., Hooijer, G.K.J. et al. (2019) A NOTCH feed-forward loop drives reprogramming from adrenergic to mesenchymal state in neuroblastoma. Nat Commun, 10, 1530.

43. Zage, P.E., Nolo, R., Fang, W., Stewart, J., Garcia-Manero, G. and Zweidler-McKay, P.A. (2012) Notch pathway activation induces neuroblastoma tumor cell growth arrest. Pediatr Blood Cancer, 58, 682–689.

44. Waldeck, K., Cullinane, C., Ardley, K., Shortt, J., Martin, B., Tothill, R.W., Li, J., Johnstone, R.W., McArthur, G.A., Hicks, R.J. et al. (2016) Long term, continuous exposure to panobinostat induces terminal differentiation and long term survival in the TH-MYCN neuroblastoma mouse model. Int J Cancer, 139, 194–204.

45. Chiu, C.Y., Kuo, K.K., Kuo, T.L., Lee, K.T. and Cheng, K.H. (2012) The activation of MEK/ERK signaling pathway by bone morphogenetic protein 4 to increase hepatocellular carcinoma cell proliferation and migration. Mol Cancer Res, 10, 415–427.

46. Lauzon, M.A. and Faucheux, N. (2018) A small peptide derived from BMP-9 can increase the effect of bFGF and NGF on SH-SY5Y cells differentiation. Mol Cell Neurosci, 88, 83–92.

