## Supplementary figures and images for "A Wnt-BMP4 signalling axis induces MSX and NOTCH proteins and promotes growth suppression and differentiation in neuroblastoma"

### Supplementary Figure 1.

**A**

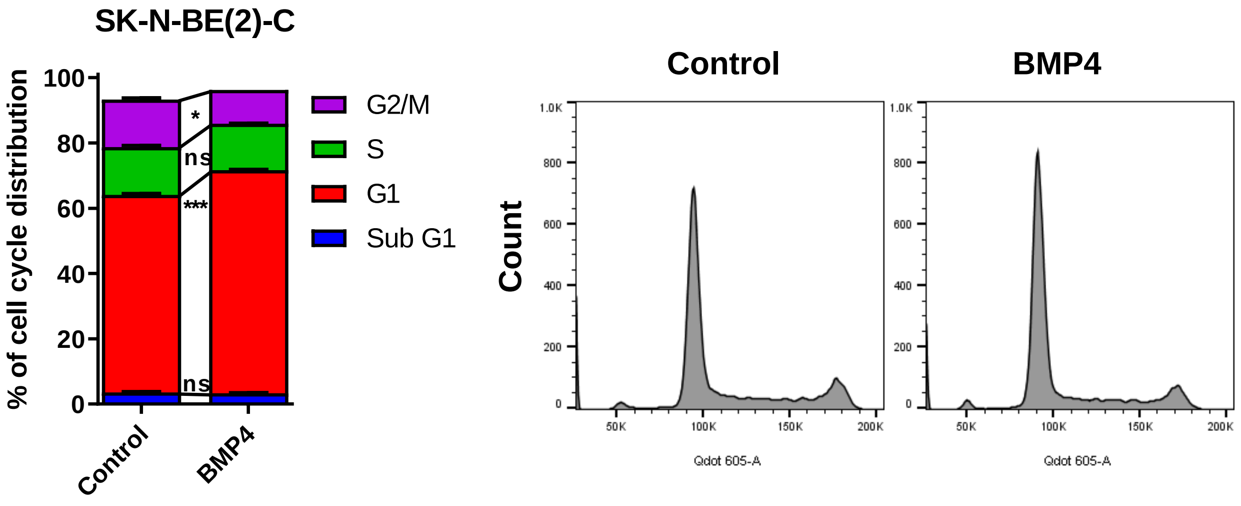


**B**

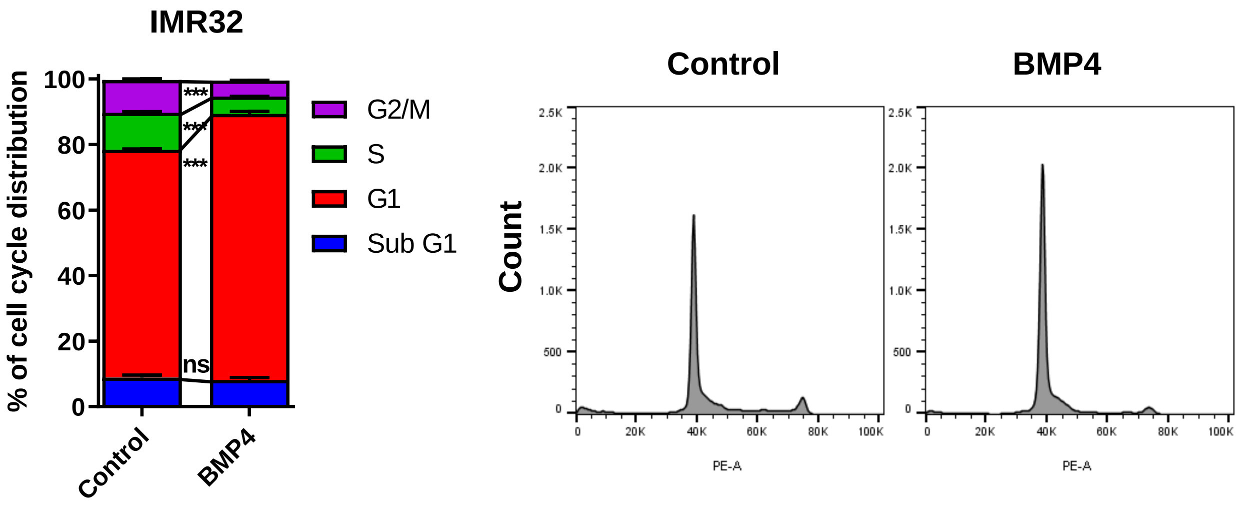


**Supplementary figure S1.**

### Supplementary Figure 2.

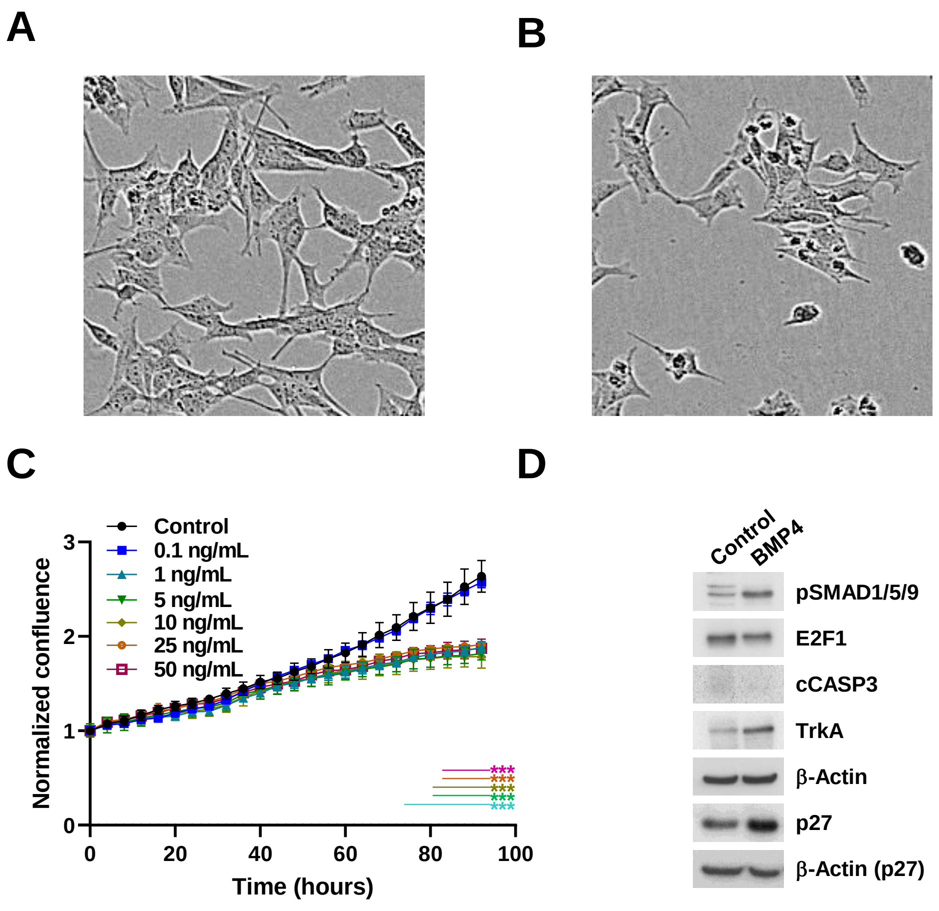


**Supplementary figure S2.**

### Supplementary Figure 3.

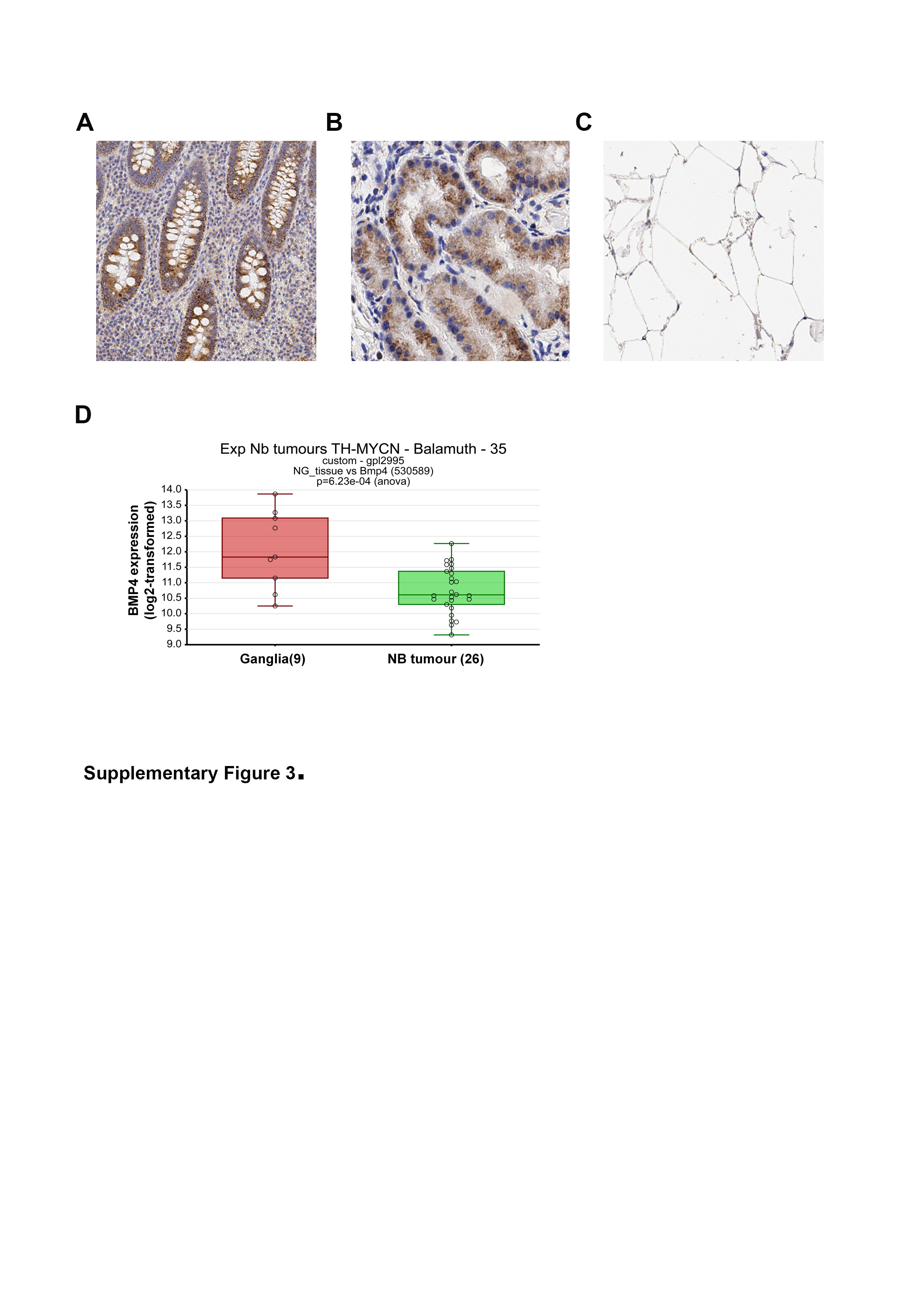

### Supplementary Figure 4.

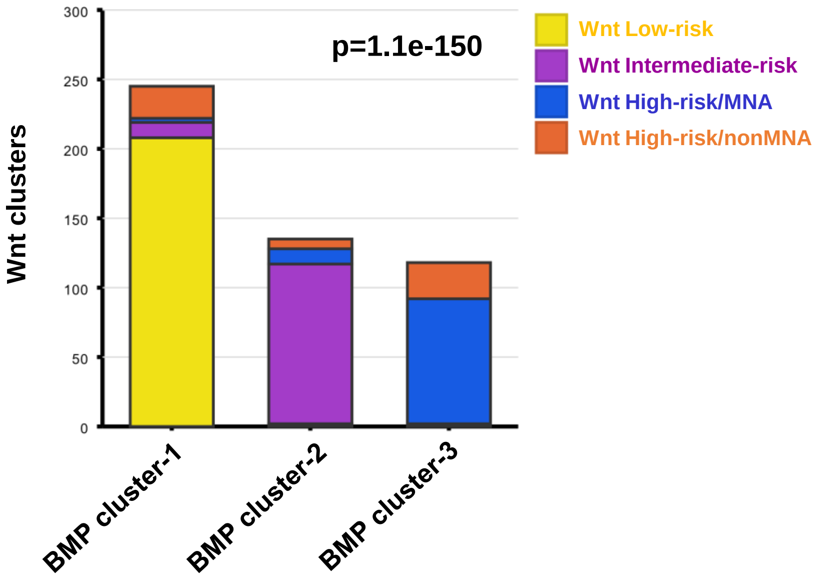


**Supplementary figure S4.**
